# The complete mitogenome of an unidentified *Oikopleura* species

**DOI:** 10.1101/2024.07.04.602139

**Authors:** Johannes Nicolaus Wibisana, Charles Plessy, Nicolas Dierckxsens, Aki Masunaga, Jiashun Miao, Nicholas M. Luscombe

## Abstract

Appendicularians are planktonic tunicates abundant all over the world. Currently, only two complete annotated mitochondrial genome assemblies are available for appendicularians, both for cryptic species of *Oikopleura dioica*. This underrepresentation of available appendicularian mitochondrial genomes limits environmental DNA sequencing (eDNA) studies that rely on mitochondrial markers as a taxonomic barcode. We report the complete mitochondrial genome assembly and annotation of an unknown appendicularian species isolated from the Amami Oshima island, Kagoshima prefecture, Japan, that has significant sequence difference with other currently available assemblies and will serve as a useful resource for ecological studies and further mitochondrial studies of appendicularians.

## 2. INTRODUCTION

Appendicularians (synonym: larvaceans) are tunicates distributed all over the world’s ocean that do not have a sessile stage, remaining free-swimming throughout its life cycle, and construct a cellulose “house” which is used for feeding and protection [1]. The best studied appendicularian is the ∼4 mm length *O. dioica* [2], but there are also large species with a body size ranging between 3–10 cm [3]. Appendicularian mitochondrial genomes use the ascidian mitochondrial genetic code [4, 5, 6, 7, 8], which differs from the invertebrate one by the reassignment of AGR codons from serine to glycine. In one clade within appendicularians containing *O. dioica*, homopolymers interrupt coding sequences and are resolved to hexamers by an unknown editing process [4, 6, 7]. In this study, we sequenced the mitochondrial genome of an unknown appendicularian species sampled from the Amami Oshima island, Japan, in order to increase the taxonomic power of eDNA studies based on the sequence of mitochondrial genes.

## 3. MATERIALS AND METHODS

### A. Sample collection and DNA extraction and sequencing

We collected specimens at Tamari harbor, Amami Oshima island, Kagoshima prefecture, Japan (28.41491667 N 129.59016667 E) in July 2023. DNA extraction was done by firstly washing samples with 5 mL of filtered autoclaved seawater 3 times before resuspending in 200 μl of lysis buffer from the MagAttract HMW DNA Kit (Qiagen, USA) with 20 μL of 10 μg/mL proteinase K and incubated for 1 h at 56°C. Next, 50 μL of 5 M NaCl was added before centrifugation of the mixture (5000 × g at 4°C) for 15 min. The supernatant was transferred into a new microtube and mixed with 400 μL of 100% EtOH and 5 μL of glycogen (20 mg/mL) and cooled at -80°C for 20 min. Further centrifugation at 6250 × g, 4°C for 5 min was performed and the supernatant removed. The obtained pellet was then washed with 1 mL of cold 70% ethanol, centrifuged, and air-dried for 5 min. The DNA was then resuspended in nuclease free water and quantified using a Qubit 3 Fluorometer (Thermo Fisher, USA). Quality control of obtained DNA was performed using Agilent 4200 TapeStation (Agilent, USA). The sequencing was performed on a PacBio Sequel II sequencer.

### B. Assembly

The sequenced reads were assembled with NOVOLoci (https://github.com/ndierckx/NOVOLoci). A partial PacBio read sequence found by a BLAST [9] search of cytochrome c oxidase subunit 1 from Okinawa *O. dioica* [7] to the raw whole DNA reads was used as a seed sequence. An initial assembly revealed two haplotypes. We assembled them separately by using new seeds extracted from the regions that contain the polymorphisms in forward and reverse direction and concatenated the assembled sequences. The coverage plot was generated by mapping the sequencing reads that were used to assemble the mitogenome to the assembly itself using minimap version 2.28-r1209 [10].

### C. Annotation

We annotated the assembly with MITOS2 v2.1.9 [11] using the ascidian mitochondrial genetic code [5]. ARWEN version 1.2.3 [12] was used in addition to annotate putative tRNAs.

### D. Phylogenetic tree

Protein-coding mitochondrial sequences were extracted from GenBank records with EMBOSS [13] and codon-aligned manually in SeaView 5.0.5 [14] after a first alignment with Clustal Omega version 1.2.4 [15]. The phylogenetic tree was computed with IQ-TREE version 2.0.7 [16] using the command-line options -T AUTO –runs 3 –polytomy –ufboot 1000 -m MFP -au -zb 1000, with one partition per gene. Sequences, accession numbers, and trees are available in the supplemental material and on Zenodo (doi:10.5281/zenodo.12660907).

### E. Protein structure prediction

3D protein structure was predicted using colabfold version 1.5.5 [17] and visualized with PyMOL [18].

## 4. RESULTS AND DISCUSSION

We noticed large appendicularians during a sampling trip targeting *O. dioica* in the Amami Oshima island, Kagoshima, Japan. We took the opportunity to collect several of these large appendicularians and sequenced a single individual, from which we assembled a circular mitogenome of length 13,058 bp (Fig. 1). We mapped the fraction of sequencing reads that were used for the assembly to the assembled sequence and obtained a sequencing depth between 36–176 × and an average depth of 122.4 × (Fig. S1).

**Fig. 1.**
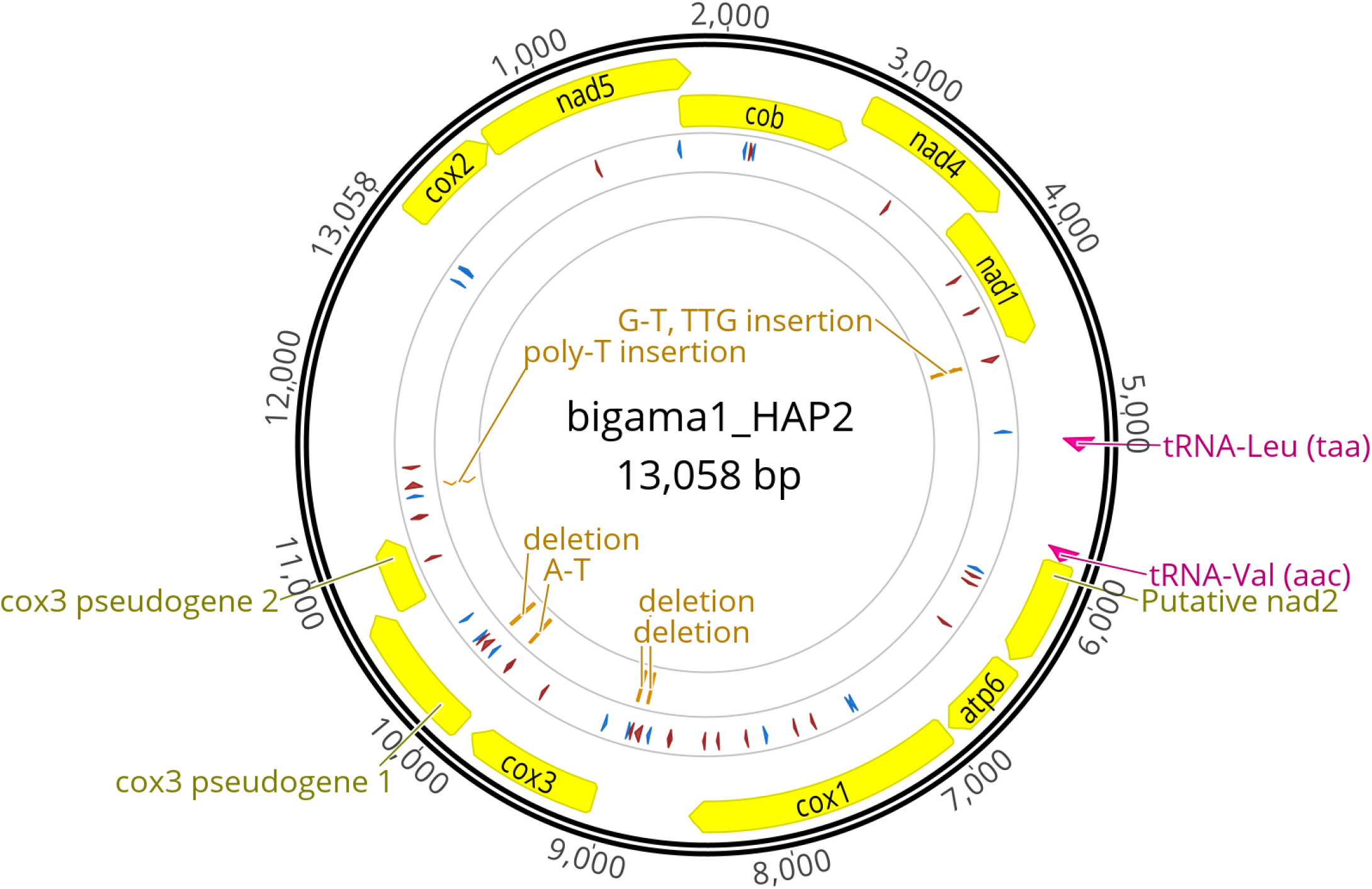
Circular plot generated by Geneious Prime version 2024.0.5. Protein coding genes and tRNAs are displayed on a yellow and pink background respectively. Unknown proteins and transcript models are displayed in gray. The outer circle illustrates the homopolymers; 6 or more successive Ts (forward) or As (reverse) in red and Cs (forward) or Gs (reverse) in blue. Abbreviations as follows, *cox2*: cytochrome c oxidase subunit 2, *nad5*: NADH dehydrogenase subunit 5, *cob*: cytochrome b, *nad4*: NADH dehydrogenase subunit 4, *nad1*: NADH dehydrogenase subunit 1, putative *nad2*: putative NADH dehydrogenase subunit 2, *atp6*: ATP synthase Fo subunit 6, *cox1*: cytochrome c oxidase subunit 1, *cox3*: cytochrome c oxidase subunit 3.

The mitogenome does not possess stretches of homopolymers like the ones observed in *O. dioica*. There is also no evidence of mitochondrial introns. Thus, we could translate its open reading frames (ORFs) with no interruptions. We found a total of 9 protein coding genes, two pseudogenes (fragments of *cox3*) and 2 tRNA genes, all on the same strand (Table 1), but could not annotate ribosomal RNA genes, although the length of the remaining unannotated regions suggest that they may be present. We were only able to detect tRNA genes for Leucine and Valine.

**Table 1.**
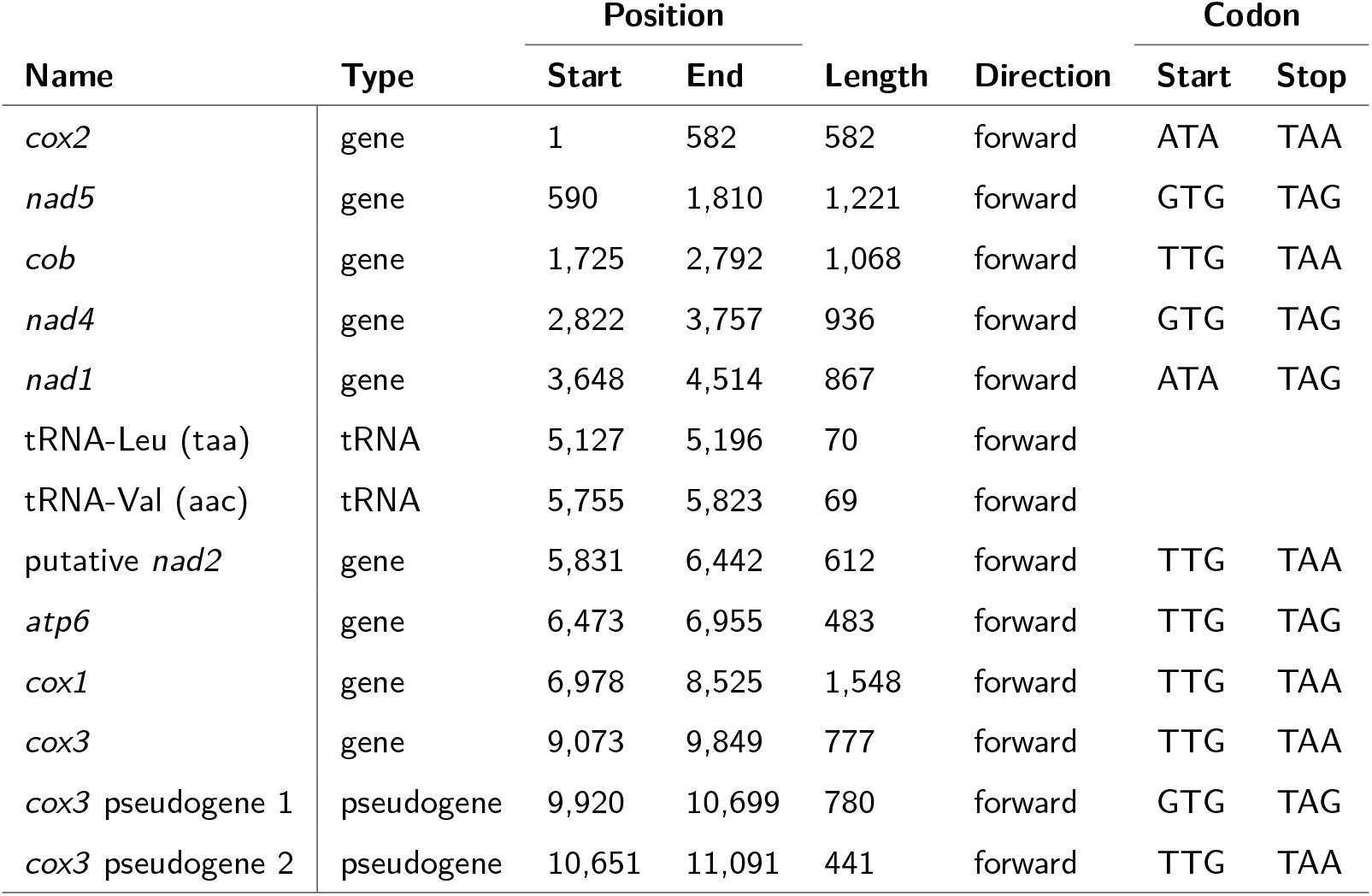
Table of genes detected including position, length, direction, start and stop codons.

| Name | Type | Position |  | Length | Direction | Codon |  |
| --- | --- | --- | --- | --- | --- | --- | --- |
|  |  | Start | End |  |  | Start | Stop |
| *cox2* | gene | 1 | 582 | 582 | forward | ATA | TAA |
| *nad5* | gene | 590 | 1,810 | 1,221 | forward | GTG | TAG |
| *cob* | gene | 1,725 | 2,792 | 1,068 | forward | TTG | TAA |
| *nad4* | gene | 2,822 | 3,757 | 936 | forward | GTG | TAG |
| *nad1* | gene | 3,648 | 4,514 | 867 | forward | ATA | TAG |
| tRNA-Leu (taa) | tRNA | 5,127 | 5,196 | 70 | forward |  |  |
| tRNA-Val (aac) | tRNA | 5,755 | 5,823 | 69 | forward |  |  |
| putative *nad2* | gene | 5,831 | 6,442 | 612 | forward | TTG | TAA |
| *atp6* | gene | 6,473 | 6,955 | 483 | forward | TTG | TAG |
| *cox1* | gene | 6,978 | 8,525 | 1,548 | forward | TTG | TAA |
| *cox3* | gene | 9,073 | 9,849 | 777 | forward | TTG | TAA |
| *cox3* pseudogene 1 | pseudogene | 9,920 | 10,699 | 780 | forward | GTG | TAG |
| *cox3* pseudogene 2 | pseudogene | 10,651 | 11,091 | 441 | forward | TTG | TAA |

Eight out of the nine protein-coding genes match known mitochondrial proteins without ambiguity. We also found an ORF with homology to the putative NADH dehydrogenase 2 (*nad2*) reported by Klirs et al. [6], and we also found matches in other appendicularian species, confirming its presence across *Oikopleuridae* (Fig. S2). Searches using BLASTp on the non-redundant protein sequences database did not yield hits, and a tBLASTx search on whole-genome shotgun contigs database of tunicates (taxid:7712) matched a predicted *Oikopleura longicauda* mitochondrial contig (SCLD01139119.1) which is different from the one we used for the phylogenetic analysis and misses some of the eight expected mitochondrial proteins. The predicted structure of the putative nad2 using colabfold [17] consists of alpha helices (Fig. S3), similar to reported nad2 protein from human (PDB IDs: 5XTC chain Q). This observation might have been caused by tunicates having fast evolving mitochondria [19], and the coverage gap in the database that currently is available.

We extended the automatic gene annotation to the longest ORF which has stop codons (TAA, TAG) and start codons (TTG, ATA, ATG, GTG) accepted by the ascidian mitochondrial code table 13. Nevertheless, due to the variability of initiation tRNA, we cannot rule out the possibility of the translation start codon being different, for example *Halocynthia roretzi* uses ATT as a start codon [20].

From the same individual we found 2 distinct mitogenomes with several single nucleotide and insertion or deletion differences, exclusively in non-coding regions (Fig. 1). We chose the mitogenome with the major variant (63% vs 37%) for the analysis performed in this study. Furthermore, these two haplotypes is supported by homopolymer variants at multiple different loci. Possible explanations for this result are heteroplasmy or that the mitogenome consists of concatemers.

The codon usage (table S1) shows that, while TGA codes for tryptophan in tunicates, it is used in less than 5% of the tryptophan positions. Furthermore, these TGA codons were only found at the most N-terminal region of *cox2*, which is not well supported by alignment to other appendicularians, and has a possible alternative start site downstream of these codons. Thus, depending on the real position of *cox2*’s translation start site, it is possible that the TGA codon is not used in this genome, similar to what was reported for *O. longicauda* on the *cox1* and *cob* genes [5]. Other than that, there are several other codon biases, such as towards TTG (39.7% and TTA (30.4%) for leucine. Another bias is present towards GTG that is coding for valine (56.7%).

The phylogenetic tree using protein-coding mitochondrial sequences (Fig. 2) shows that this unknown species belongs to the clade of larvaceans that include *Bathochordaeus, Mesochordaeus* and *Oikopleura longicauda* but not *O. dioica*. This clade was was also found in a phylogenetic analysis of ribosomal protein sequences by Naville *et al*., 2019 [21]. The split between *O. dioica* and the other appendicularians in our tree corresponds to the bioluminescent / non-bioluminescent classification of Galt *et al*., 1985 [22]. This is also reflected in the situation of homopolymers which are not abundant in this mitogenome, similar to *O. longicauda* [21] and not *O. dioica* [7]. As the *Oikopleura* genus is polyphylic in our phylogenetic anaylysis and that of others, further work not in the scope of this manuscript will be needed to resolve which genus has to be corrected.

**Fig. 2.**
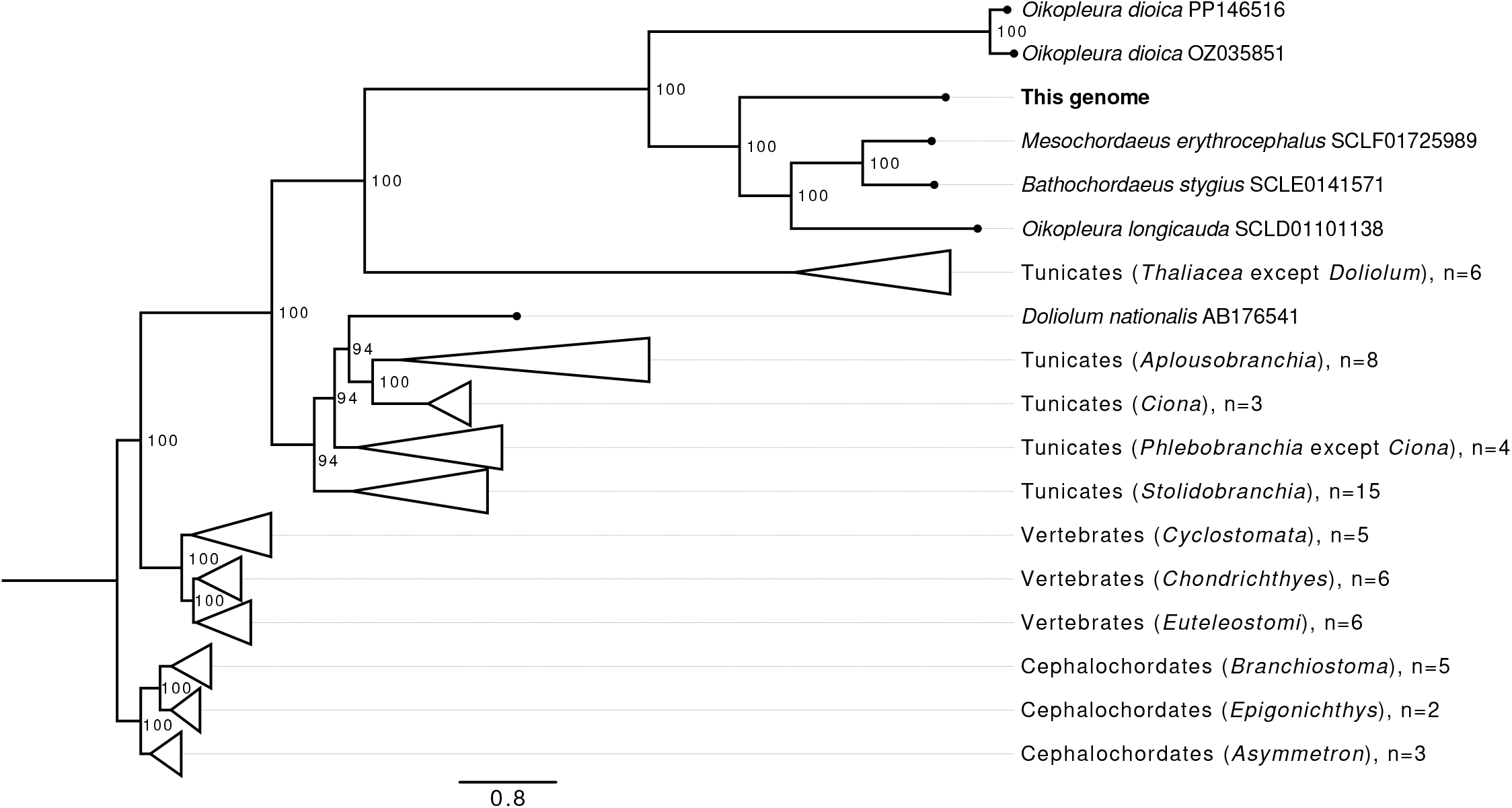
Phylogenetic tree computed on codon-aligned mitochondrial genome protein-coding genes collected from publicly available aquatic chordate genomes.ter

## 5. DISCUSSION AND CONCLUSION

We present here the complete mitogenome of an unidentified *Oikopleura* species. While the assembly is circular, it is possible that the genome is linear concatenate or is present in multiple copies in a circle due to the length of the control region. This unidentified species seems to have a mitogenome that closer resembles that of the lineage of *O. longicauda* instead of *O. dioica*. We project that the data produced in this study will be useful in future eDNA studies.

## 6. ETHICAL APPROVAL

No ethical issues were involved in this study.

## 7. AUTHOR CONTRIBUTIONS

AM and JM collected samples and AM performed sequencing. JNW, ND, and CP performed bioinformatics analysis. JNW drafted the manuscript. CP, ND, and NML critically revised the manuscript. All authors approved the final manuscript and agreed to be accountable for all aspects of this work.

## 9. DATA AVAILABILITY STATEMENT

The mitochondrial genome sequence was deposited in GenBank under the accession number LC830956. The annotation and the sequences used to compute the phylogenetic tree in Fig. 2 are available in Zenodo (doi:10.5281/zenodo.12660907).

## 10. ACKNOWLEDGEMENTS

We thank Dr. Michael Mansfield for the invaluable suggestions and guidance in the construction of the phylogenetic tree. We thank the DNA Sequencing Section and the Scientific Computing and Data Analysis Section of the Research Support Division at OIST for their support. This work was supported by OIST core funding and JSPS KAKENHI grant number 23K14236.

---

**Figure S1.**
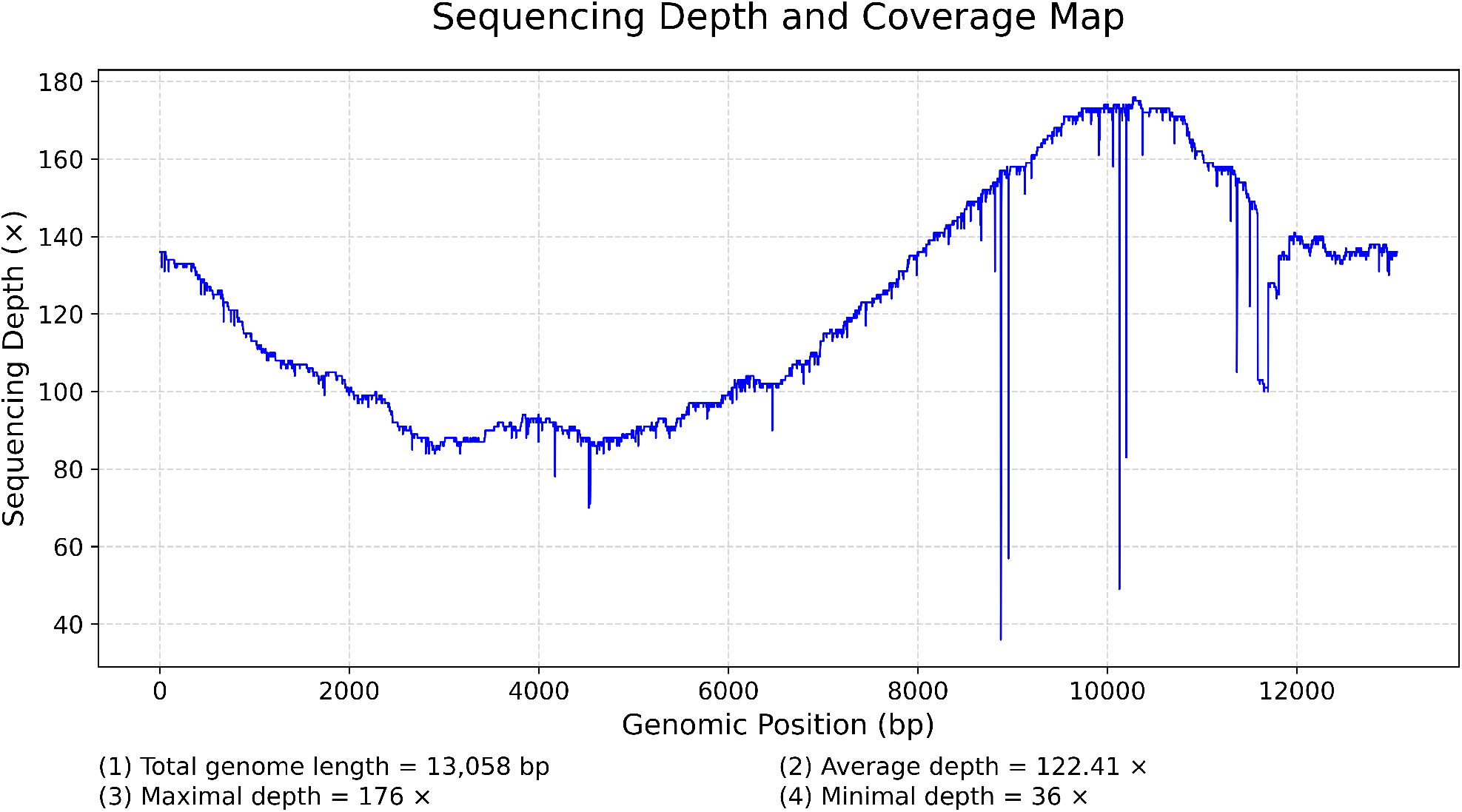
Coverage plot of the mapping of assembled sequencing reads.

**Figure S2.**
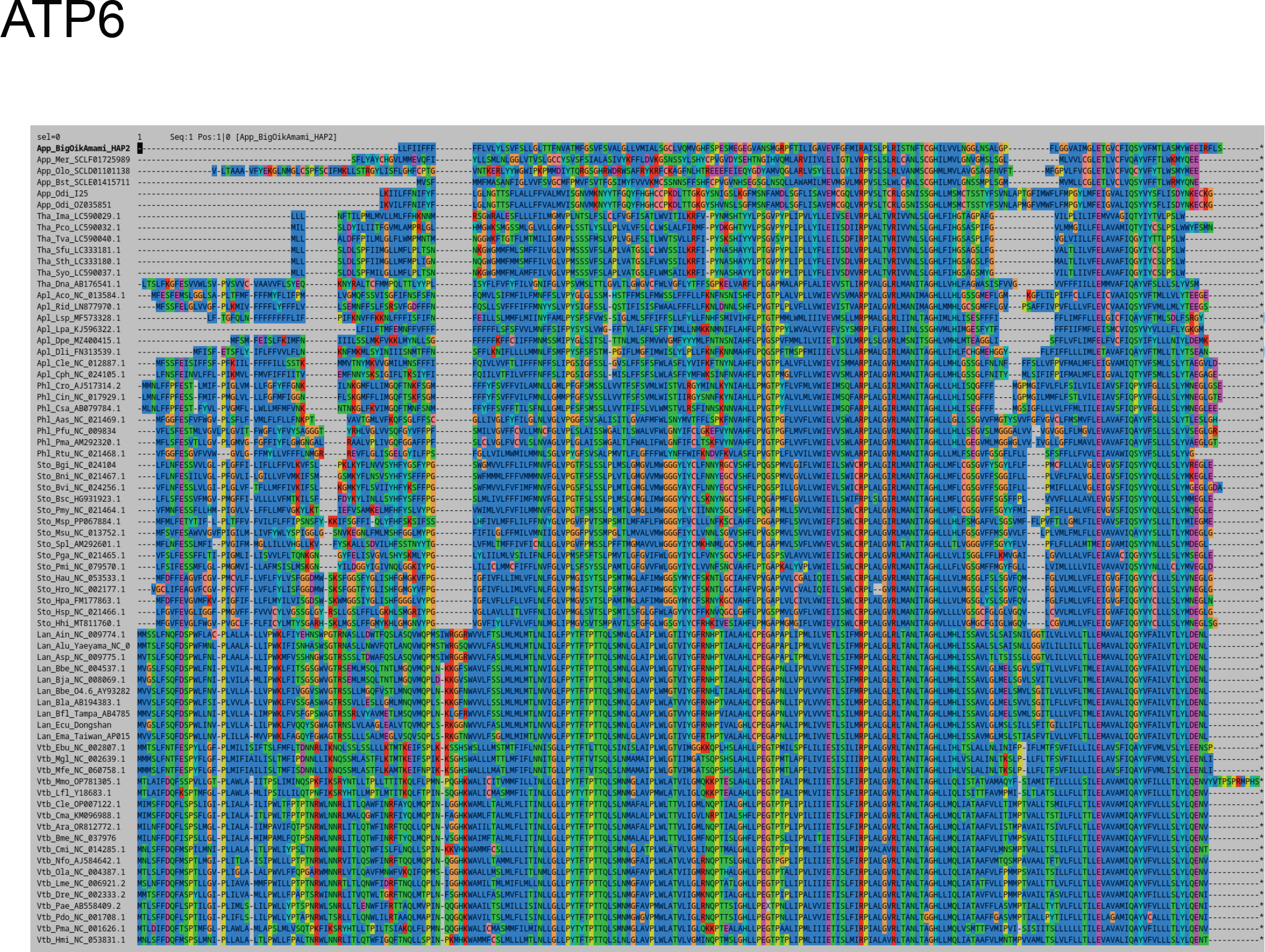
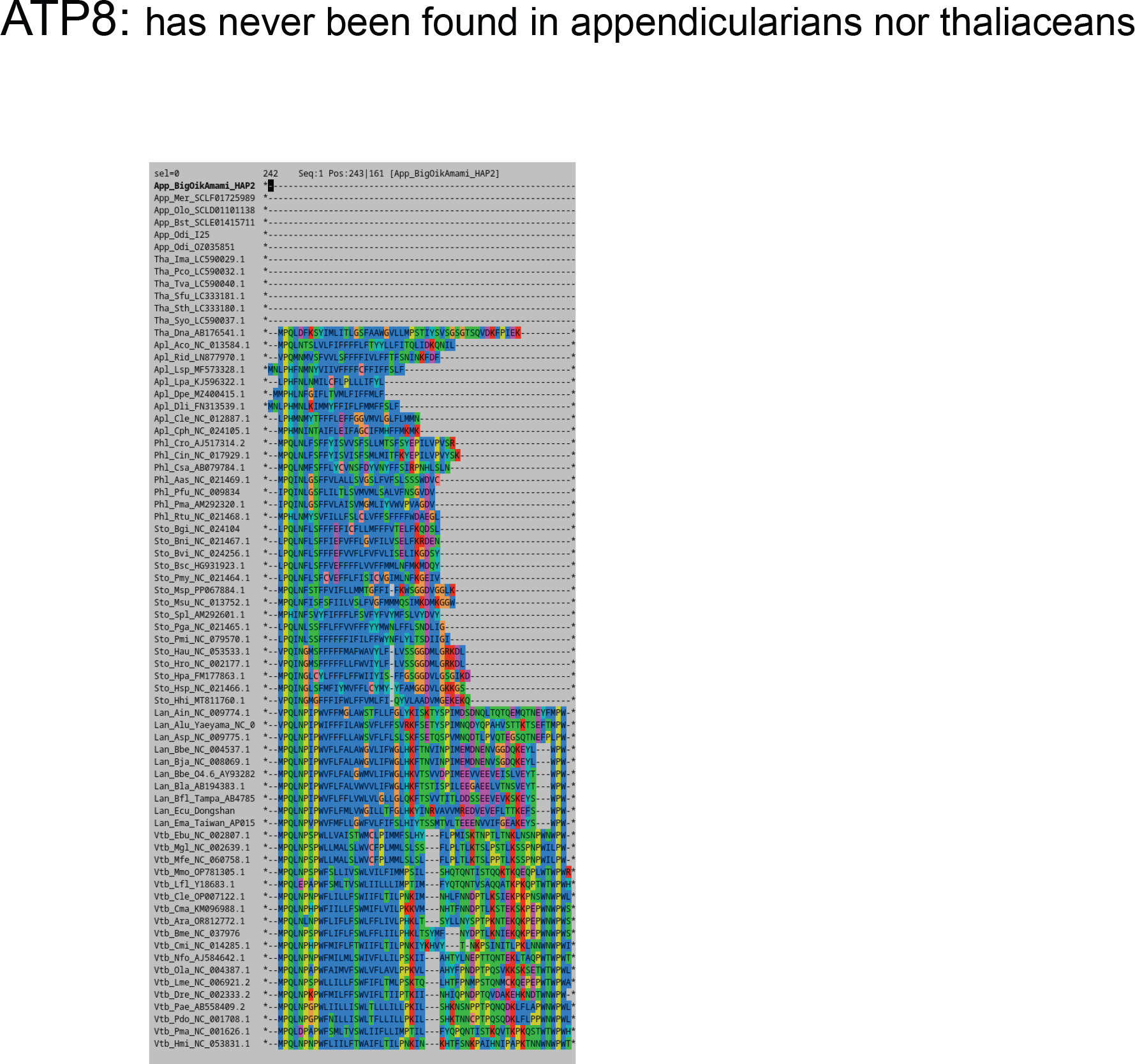
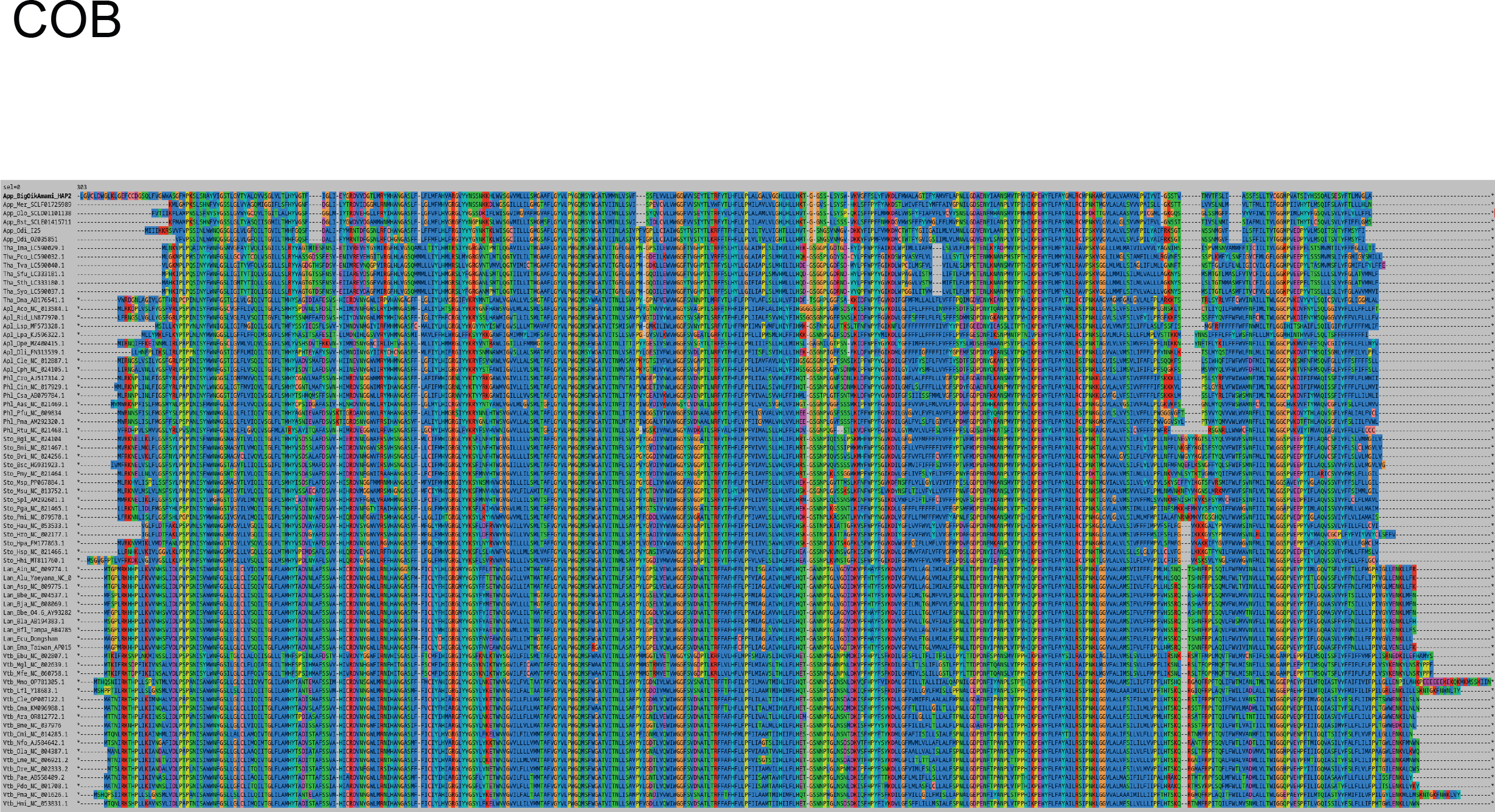
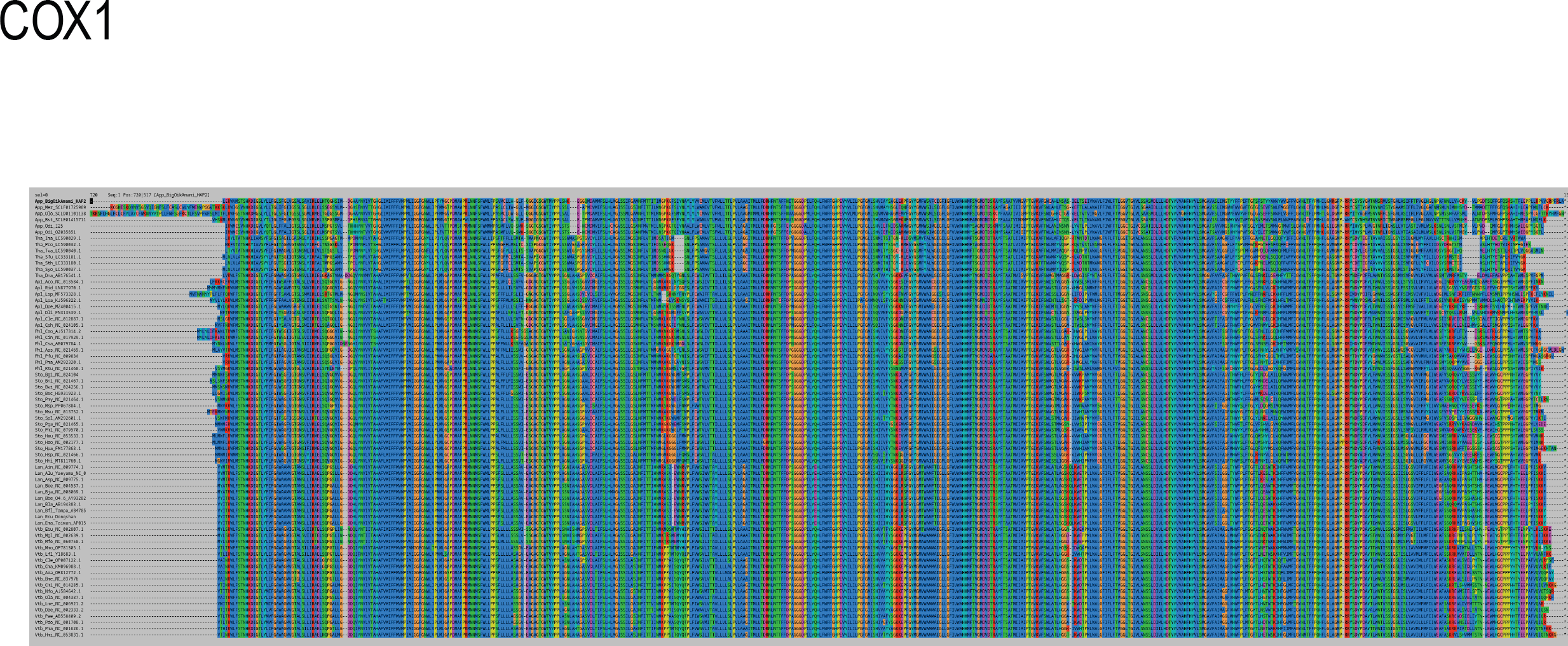
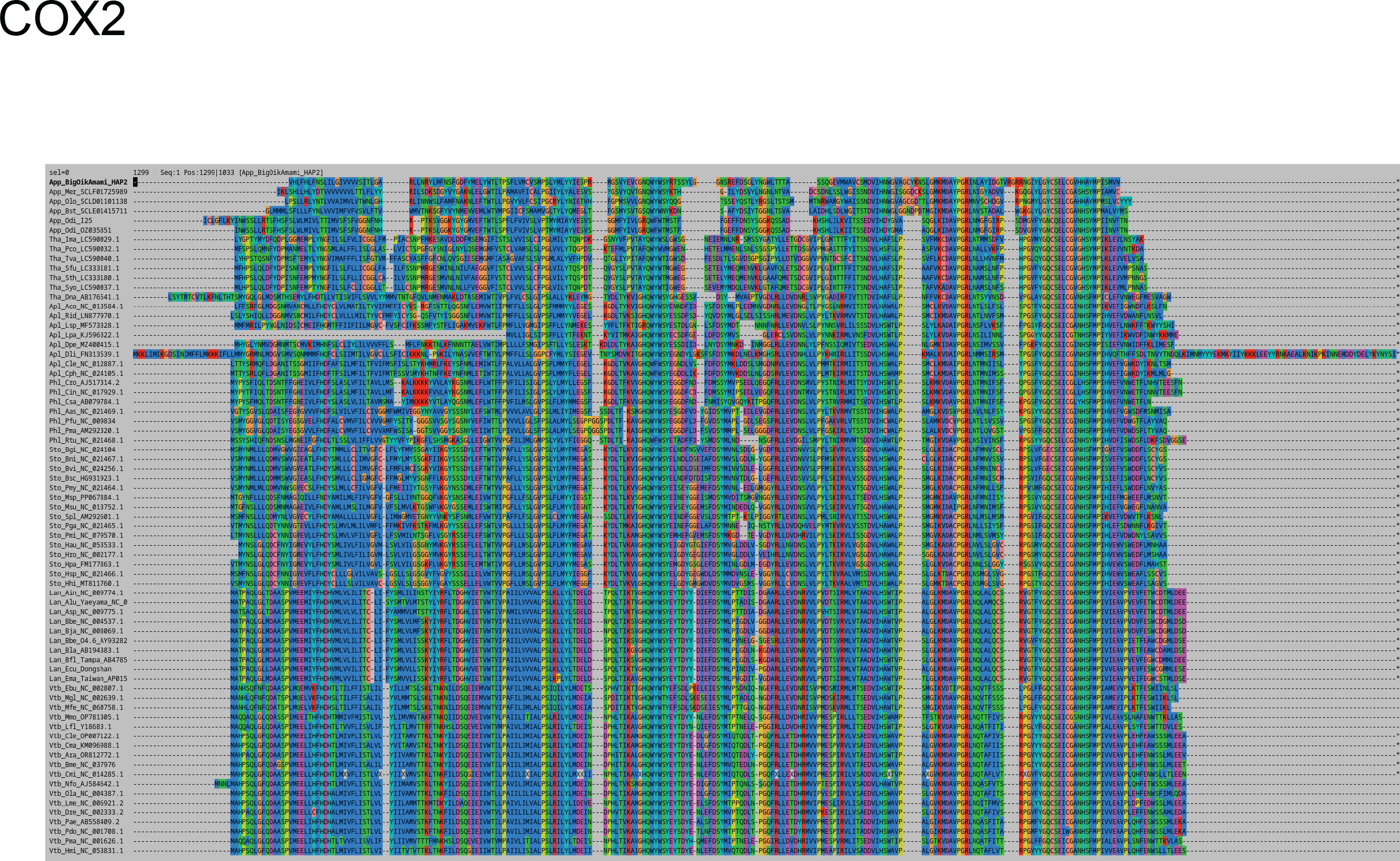
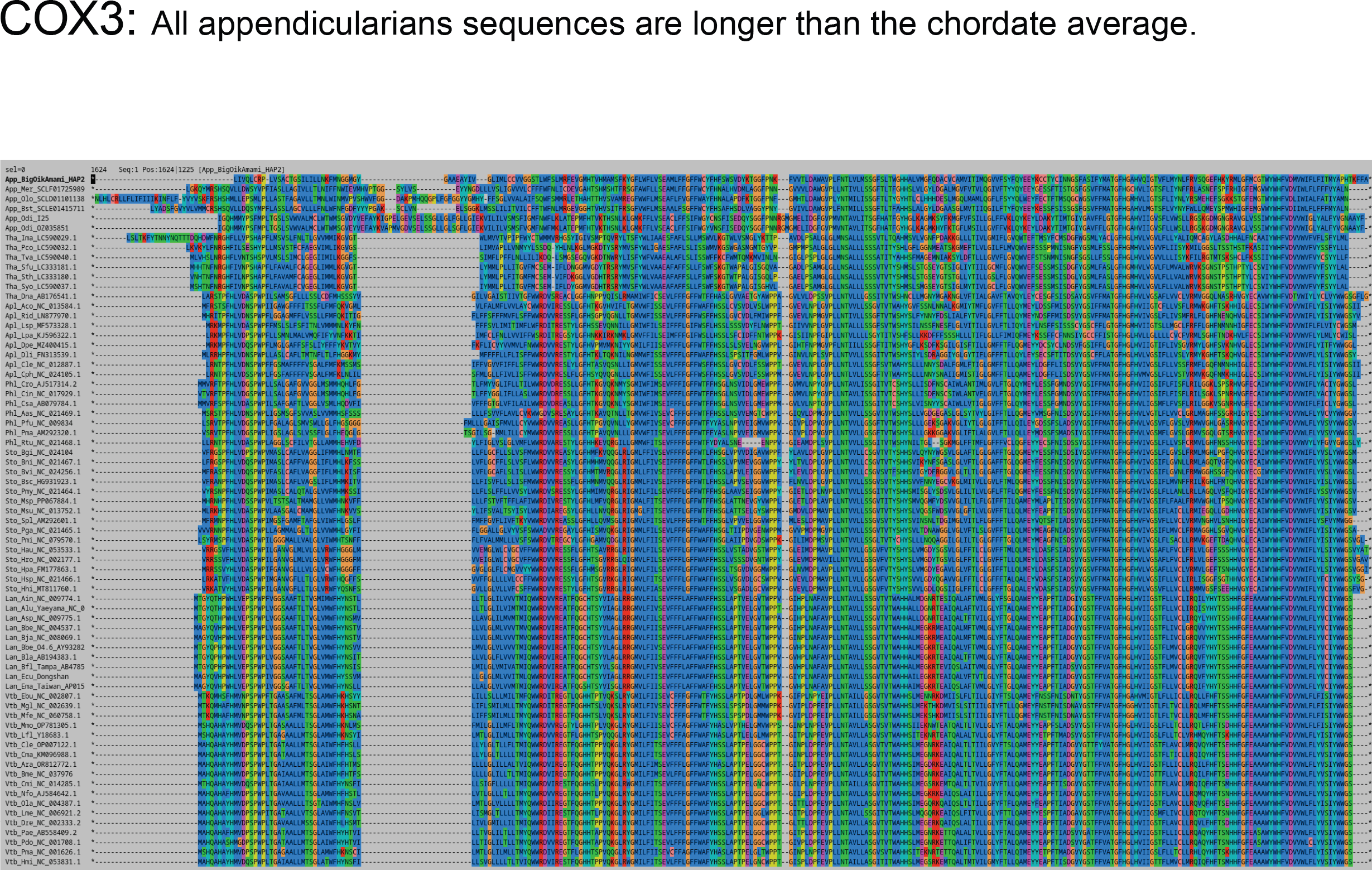
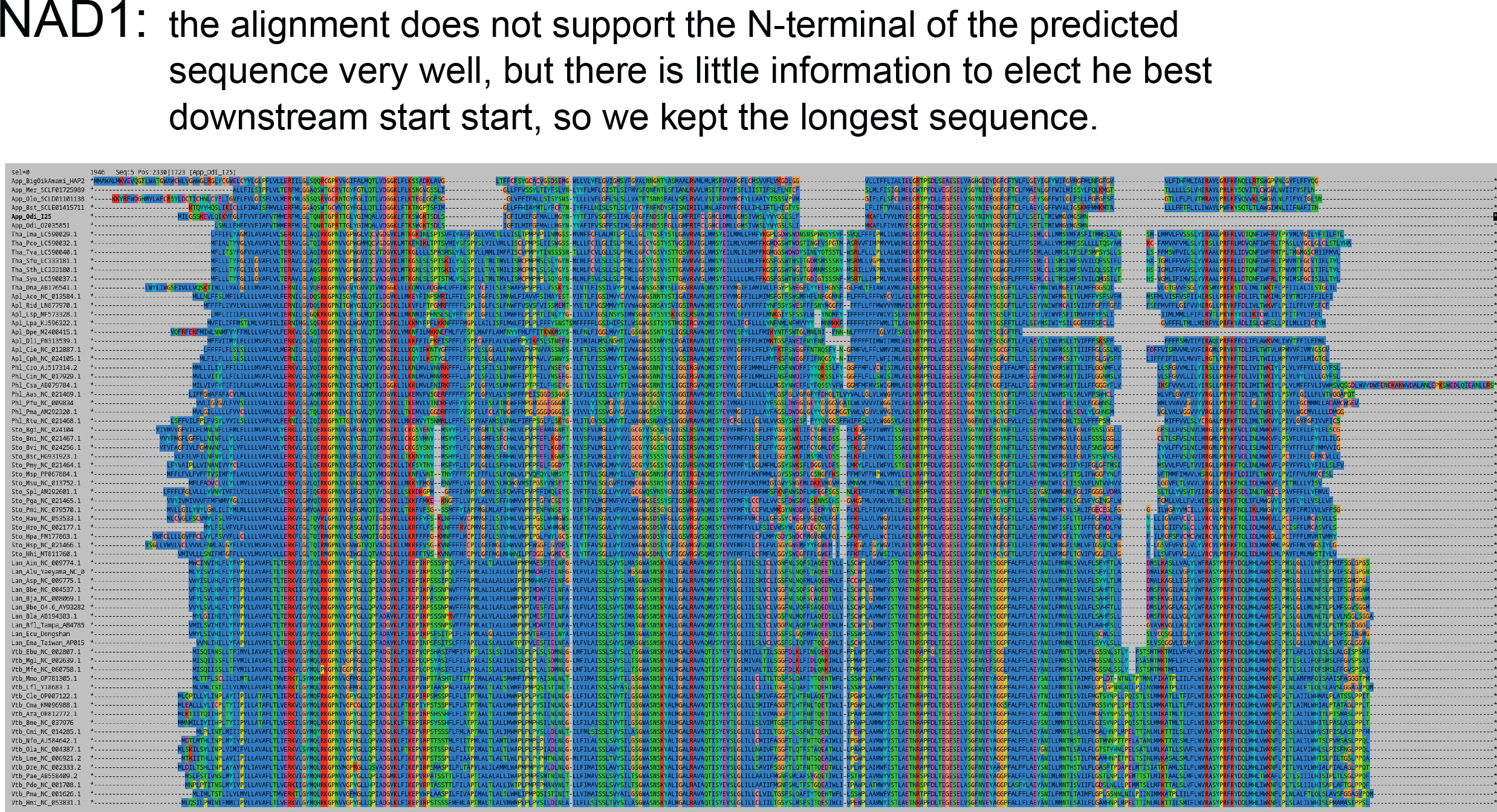
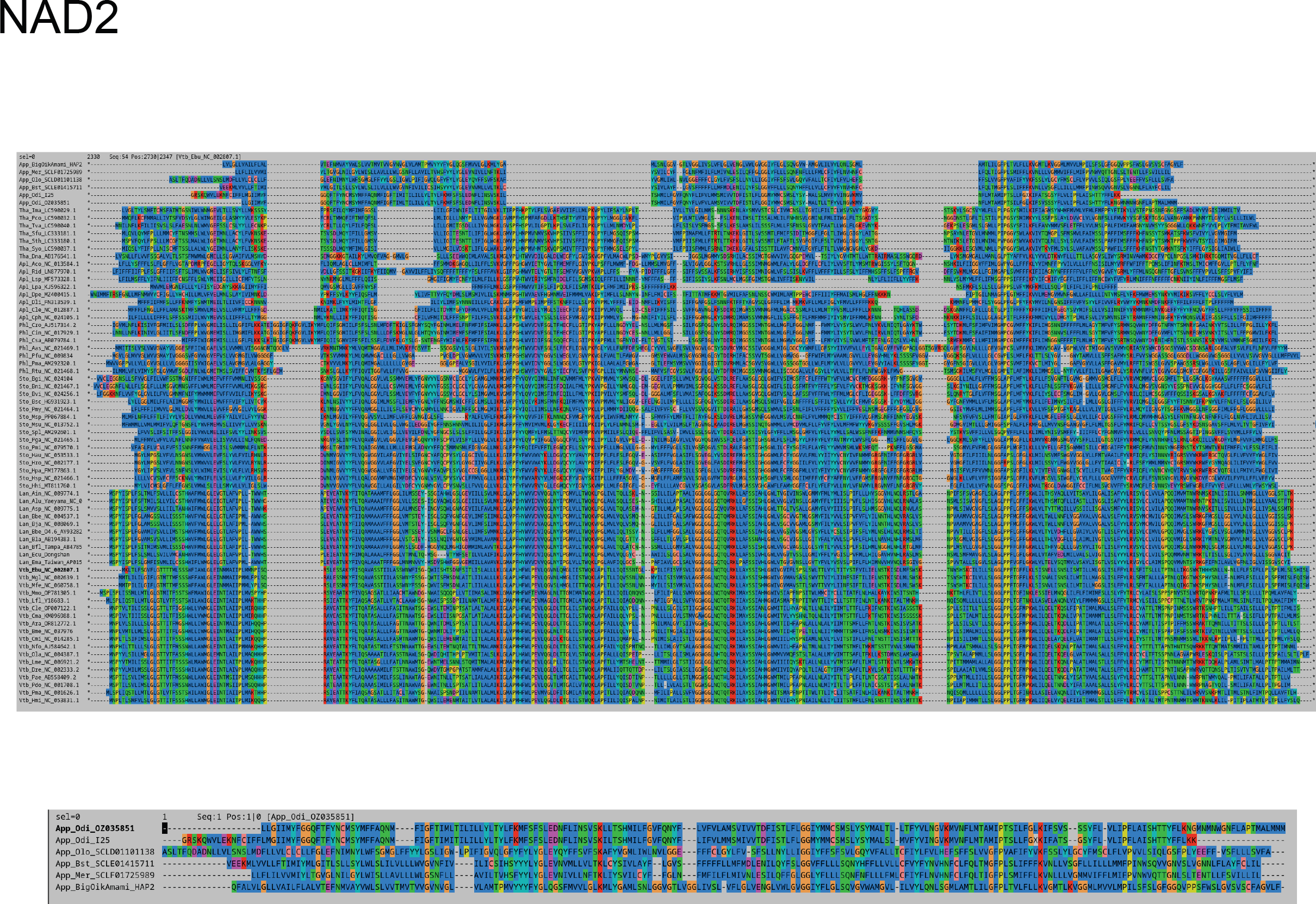
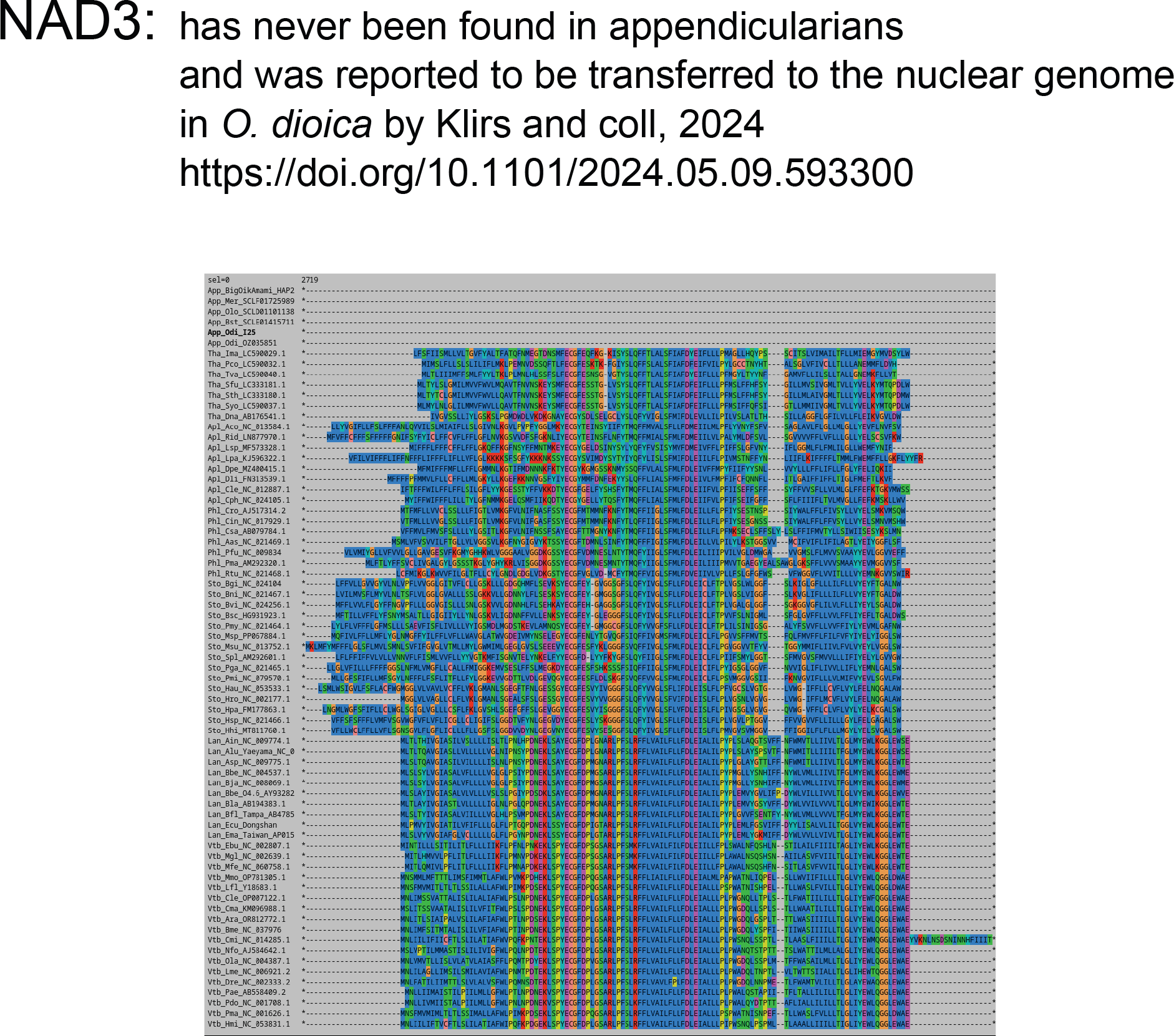
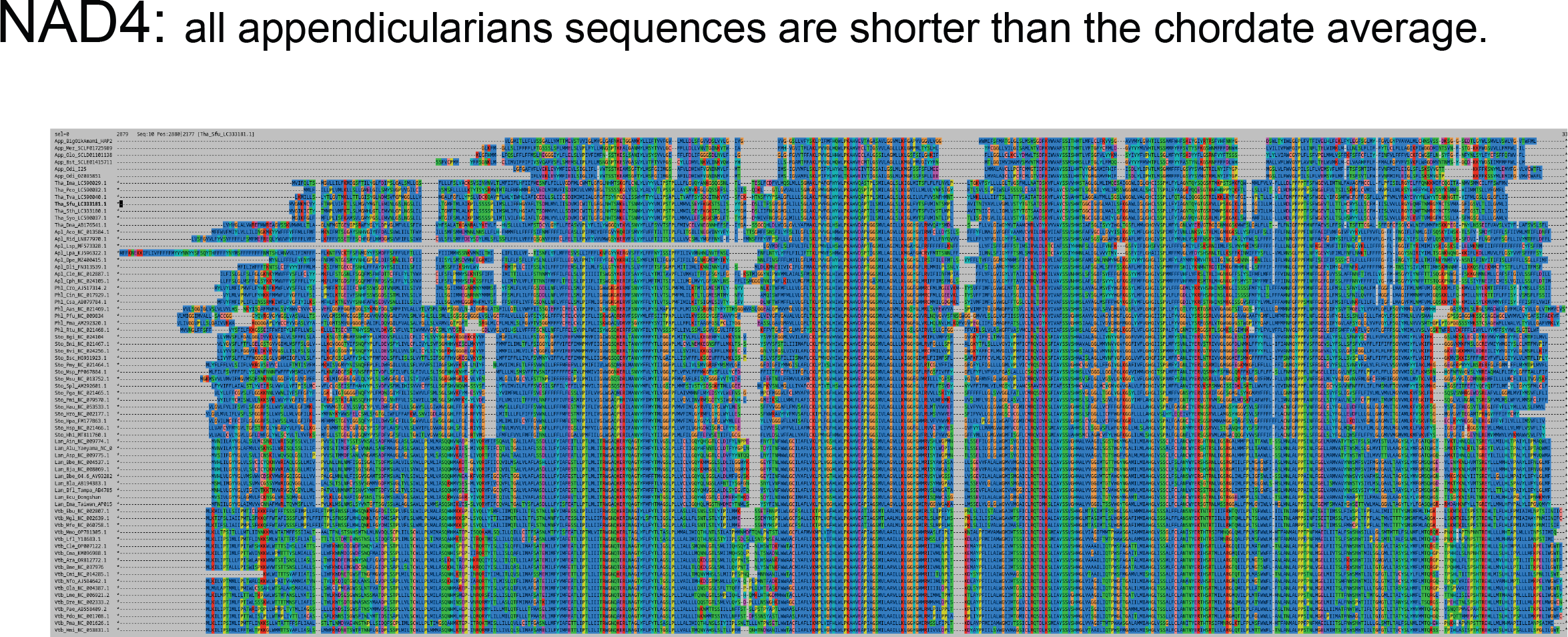
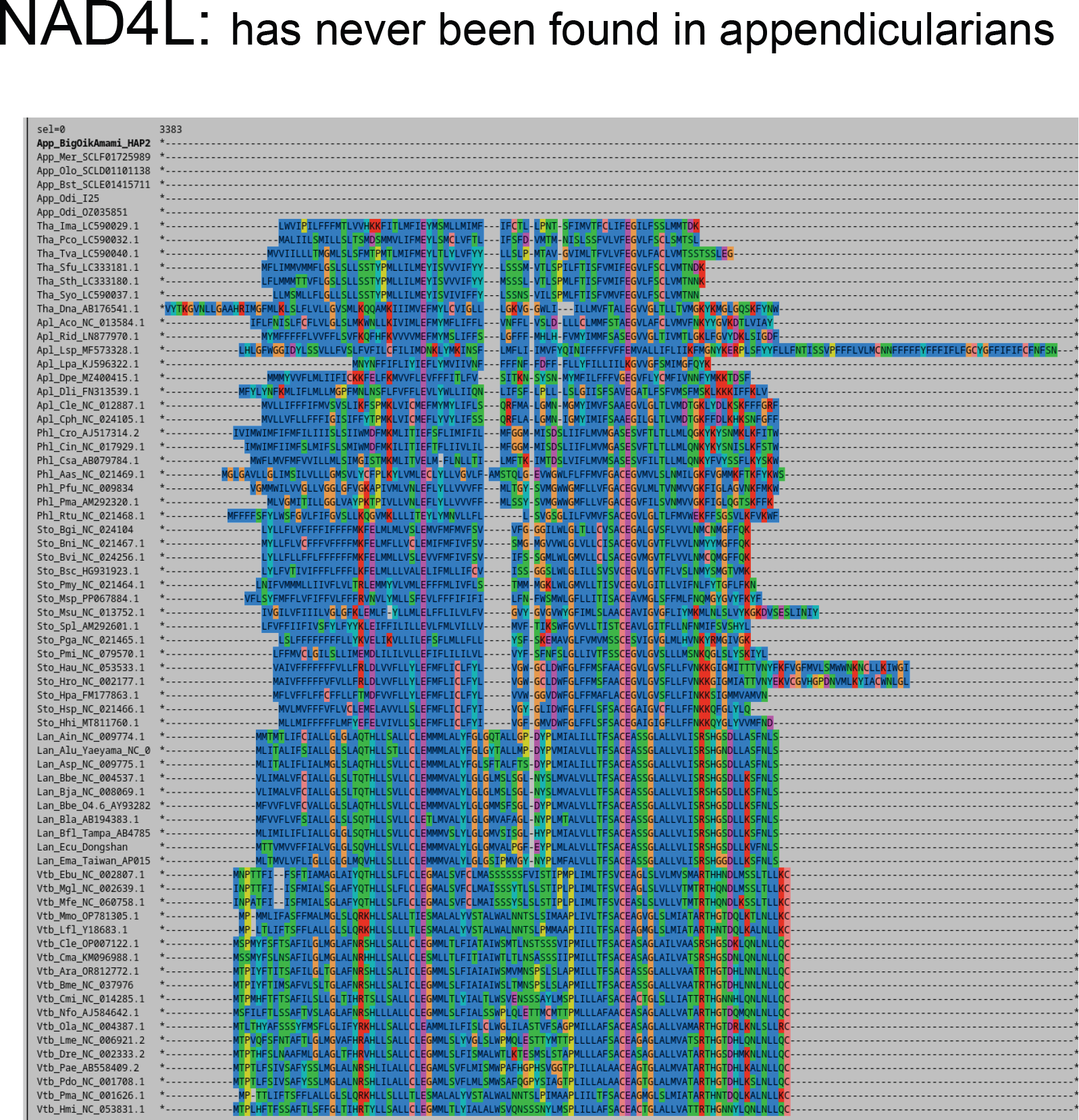
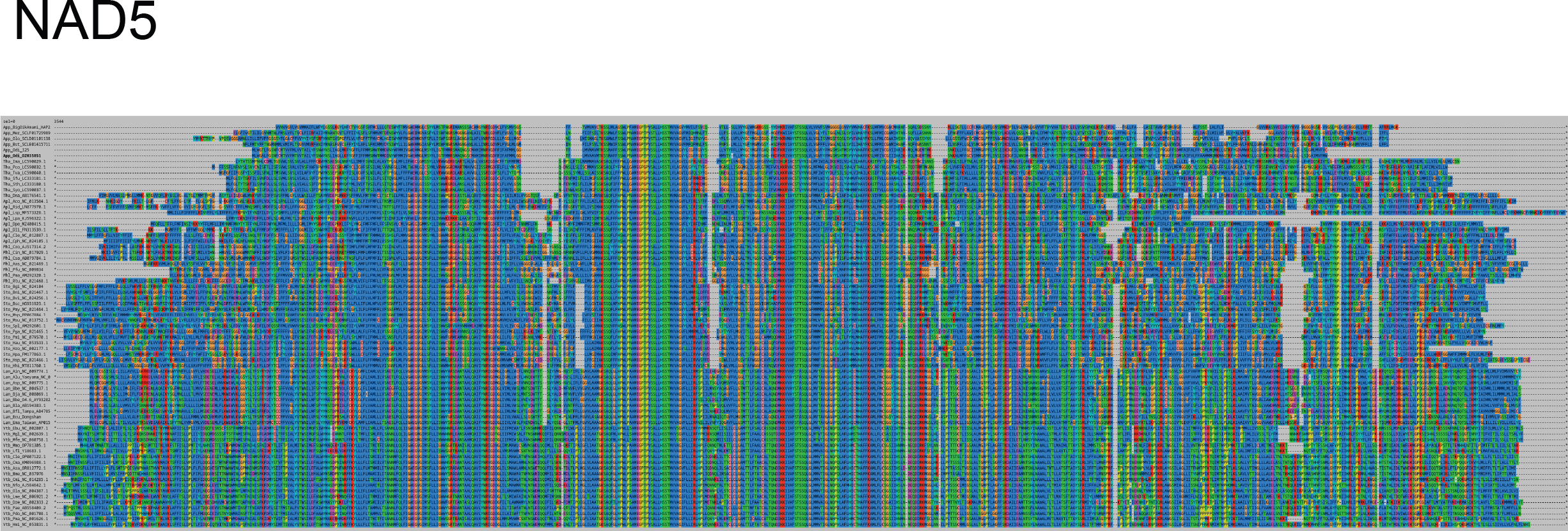
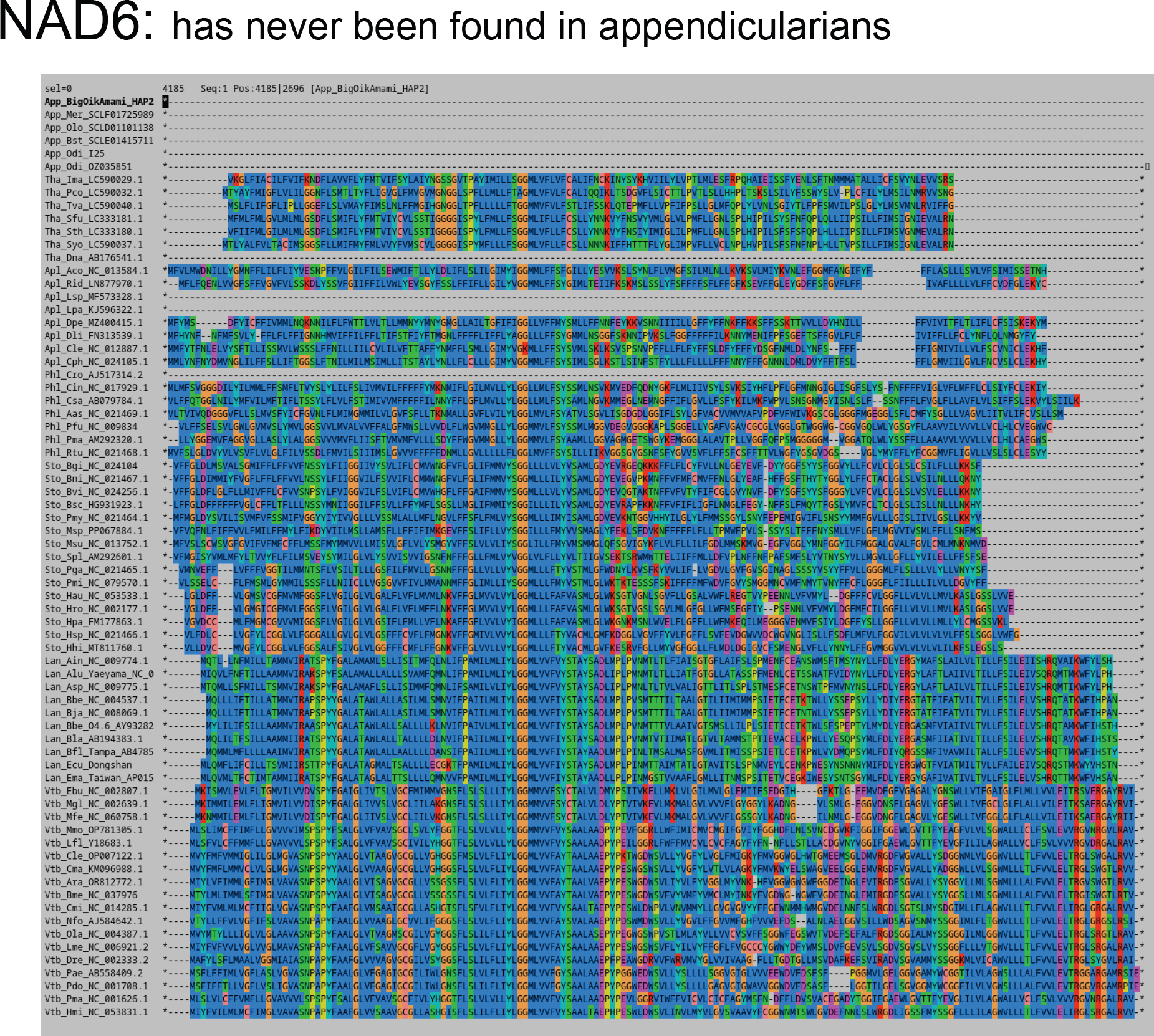
Manual alignment of protein-coding genes to other tunicates

**Figure S3.**
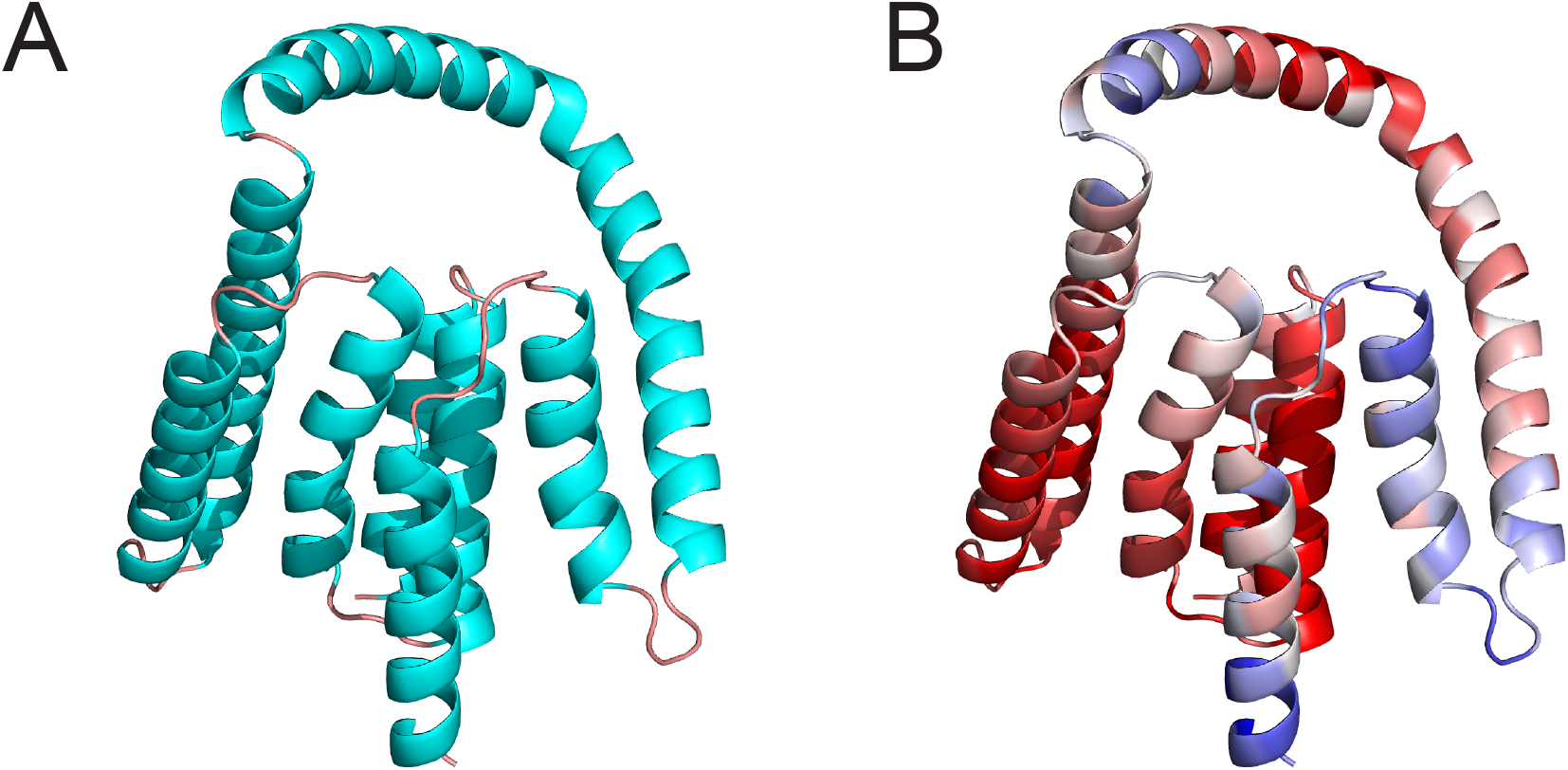
Predicted 3D structure of the putative nad2 protein. (A) Colored by secondary structure and (B) colored by pLDDT score from 30 (blue) to 60 (red).

**Table S1.** Codon usage table of protein-coding genes in this genome.

| Codon | AA | %of AA | Freq |
| --- | --- | --- | --- |
| GCA | A | 17.5% | 27 |
| GCC | A | 16.9% | 26 |
| GCG | A | 20.1% | 31 |
| GCT | A | 45.5% | 70 |
| TGC | C | 39.7% | 23 |
| TGT | C | 60.3% | 35 |
| GAC | D | 31.9% | 15 |
| GAT | D | 68.1% | 32 |
| GAA | E | 13.8% | 8 |
| GAG | E | 86.2% | 50 |
| TTC | F | 32.0% | 74 |
| TTT | F | 68.0% | 157 |
| AGA | G | 17.2% | 52 |
| AGG | G | 24.5% | 74 |
| GGA | G | 11.9% | 36 |
| GGC | G | 5.3% | 16 |
| GGG | G | 27.2% | 82 |
| GGT | G | 13.9% | 42 |
| CAC | H | 38.3% | 23 |
| CAT | H | 61.7% | 37 |
| ATC | I | 33.0% | 35 |
| ATT | I | 67.0% | 71 |
| AAA | K | 33.3% | 19 |
| AAG | K | 66.7% | 38 |
| CTA | L | 10.7% | 40 |
| CTC | L | 1.3% | 5 |
| CTG | L | 14.4% | 54 |
| CTT | L | 3.5% | 13 |
| TTA | L | 30.4% | 114 |
| TTG | L | 39.7% | 149 |
| ATA | M | 29.6% | 48 |
| ATG | M | 70.4% | 114 |
| GTG | M | 1.20% | 2 |
| TTG | M | 2.90% | 5 |
| AAC | N | 38.2% | 26 |
| AAT | N | 61.8% | 42 |
| CCA | P | 17.5% | 11 |
| CCC | P | 11.1% | 7 |
| CCG | P | 19.0% | 12 |
| CCT | P | 52.4% | 33 |
| CAA | Q | 33.3% | 12 |
| CAG | Q | 66.7% | 24 |
| CGA | R | 17.3% | 9 |
| CGC | R | 5.8% | 3 |
| CGG | R | 63.5% | 33 |
| CGT | R | 13.5% | 7 |
| AGC | S | 15.1% | 33 |
| AGT | S | 22.0% | 48 |
| TCA | S | 15.6% | 34 |
| TCC | S | 7.8% | 17 |
| TCG | S | 20.2% | 44 |
| TCT | S | 19.3% | 42 |
| ACA | T | 19.2% | 24 |
| ACC | T | 18.4% | 23 |
| ACG | T | 25.6% | 32 |
| ACT | T | 36.8% | 46 |
| GTA | V | 14.0% | 43 |
| GTC | V | 9.4% | 29 |
| GTG | V | 56.7% | 174 |
| GTT | V | 19.9% | 61 |
| TGA | W | 4.1% | 3 |
| TGG | W | 95.9% | 71 |
| TAC | Y | 27.6% | 35 |
| TAT | Y | 72.4% | 92 |
| TAA | * | 55.6% | 5 |
| TAG | * | 44.4% | 4 |

